# A low-resource reliable pipeline to democratize multi-modal connectome estimation and analysis

**DOI:** 10.1101/2021.11.01.466686

**Authors:** Jaewon Chung, Ross Lawrence, Alex Loftus, Derek Pisner, Gregory Kiar, Eric W. Bridgeford, William Gray Roncal, Consortium for Reliability and Reproducibility (CoRR), Vikram Chandrashekhar, Disa Mhembere, Sephira Ryman, Xi-Nian Zuo, Daniel S. Margulies, R. Cameron Craddock, Carey E. Priebe, Rex Jung, Vince D. Calhoun, Brian Caffo, Randal Burns, Michael P. Milham, Joshua T. Vogelstein

## Abstract

Connectomics—the study of brain networks—provides a unique and valuable opportunity to study the brain. Research in human connectomics, leveraging functional and diffusion Magnetic Resonance Imaging (MRI), is a resource-intensive practice. Typical analysis routines require significant computational capabilities and subject matter expertise. Establishing a pipeline that is low-resource, easy to use, and off-the-shelf (can be applied across multifarious datasets without parameter tuning to reliably estimate plausible connectomes), would significantly lower the barrier to entry into connectomics, thereby democratizing the field by empowering a more diverse and inclusive community of connectomists. We therefore introduce ‘MRI to Graphs’ (`m2g`). To illustrate its properties, we used `m2g` to process MRI data from 35 different studies (≈ 6,000 scans) from 15 sites without any manual intervention or parameter tuning. Every single scan yielded an estimated connectome that adhered to established properties, such as stronger ipsilateral than contralateral connections in structural connectomes, and stronger homotopic than heterotopic correlations in functional connectomes. Moreover, the connectomes estimated by `m2g` are more similar within individuals than between them, suggesting that `m2g` preserves biological variability. `m2g` is portable, and can run on a single CPU with 16 GB of RAM in less than a couple hours, or be deployed on the cloud using its docker container. All code is available on https://github.com/neurodata/m2g and documentation is available on docs.neurodata.io/m2g.

## 1 Introduction

Human brain imaging, especially Magnetic Resonance Imaging (MRI), has become a vital tool in both basic and clinical brain science [1], with new analysis techniques being developed frequently. One group of techniques concerns itself with estimation and analysis of connectomes from MRI data, allowing for the use of graph theoretic mathematics and statistics to discern both functional and structural relationships. A connectome is a comprehensive map of relationships present in the brain at a given scale and time. The estimation of a connectome revolves around the number of connections, or edges, between different areas of the brain called regions of interest (ROIs). Where the boundaries lie for each of these regions of interest depends on the parcellation method used, as there are many different ways to group regions of the brain. Edges can represent a number of relationships between ROIs. For estimated structural connectomes, which are generated using diffusion-weighted MRI scans, edges represent the quantity of white-matter tracks that connect ROIs. For functional connectomes, which are estimated from blood oxygenation level dependent (BOLD) images, edges represent the correlation in activation between pairs of regions, as neuronal firing is shortly followed by a depletion of oxygen in surrounding capillaries as neurons prepare to fire again.

The process of estimating a connectome from MRI data involves multiple steps, each with different inputs and outputs, and can be performed with a variety of parameters. There are two main approaches that can be taken in the design of a pipeline for estimating connectomes: (1) optimize the pipeline for each dataset you wish to analyze or (2) design a “general” pipeline that can be used on a variety of data. While creating an optimized pipeline for each dataset results in the “most accurate” connectomes for the data, it also limits the ability for the pipeline to be used with new data and for comparisons between datasets. Alternatively, the creation of a “standard” pipeline which can adequately estimate connectomes from a wide collection of datasets will allow for more confident comparison at the expense of potentially sub-optimal connectome estimation. While there exist multiple pipelines which perform part or all of the processes required to estimate a connectome from MRI data, few exist which are easily accessible for individuals new to the field. Both `fmriprep` [2] and `dmriprep` [3] do not estimate a connectome from provided data: rather, they serve to preprocess data for further analysis by other software. Other MRI connectome estimation pipelines often require significant background knowledge or computational resources in order to be utilized effectively. For example, while the Human Connectome Pipeline (HCP) [4] does have a docker container allowing for easy use, their structural connectome generation can take over 24 hours and require up to 24 GB while their functional connectome generation can take over 36 hours and require up to 24 GB depending on the duration of the fMRI scan.

In hopes to expand the study of connectomes to a larger group of individuals, we developed a pipeline, called “MRI to Graphs” (`m2g`), which utilizes accepted approaches for connectome generation along with multiple open-source tools. The `m2g` pipeline serves to streamline the process of estimating connectomes for both functional and structural MRI data through handling the required preprocessing, graph generation, and quality assurance from one function call. Rather than providing bespoke analyses for each unique dataset, limiting their generalizability and likely increasing computational cost, `m2g` harmonizes processing to produce highly reliable data-derived quantities across a wide variety of datasets. To further the goal of a pipeline accessible to everyone, `m2g` also provides extensive quality assurance at each data processing stage in the form of easy to understand images, using widely accepted visualization formats.

During development and testing, we used `m2g` to process 13 diffusion-weighted MRI (dMRI) studies comprising ≈720 individuals with ≈1,400 scans, and 30 functional MRI (fMRI) studies comprising ≈1,400 individuals with ≈3,500 scans—each of which was estimated using 35 different parcellation methods from the neuroparc repository [5]. Estimation of each of these connectomes was performed with one command line call, 3 CPUs, and ≈ 16 GB of RAM, in under an hour and a half from raw data to connectome. Due to the robustness of `m2g`, each connectome was estimated without issue and passed our surrogate metrics for plausibility. These connectomes, in addition to code and other data derivatives, are publicly available at https://neurodata.io/mri/.

## 2 Results

### 2.1 The m2g Pipeline

The `m2g` pipeline consists of two separate sub-pipelines: `m2g-d`, which processes diffusion-weighted MRI scans to estimate structural connectomes, and `m2g-f`, which processes BOLD functional MRI scans to estimate functional connectomes. While the `m2g-d` sub-pipeline is locally contained in `m2g`, `m2g-f` has been integrated into CPAC [6] as a Nipype pipeline, [7] which is called during the processing of functional MRI data. While the code itself was developed by our group, and the performance is monitored for consistency between updates, the `m2g-f` sub-pipeline is hosted and maintained by the FCP-INDI organization. The steps involved in each of these sub-pipelines are further discussed in the Methods section. Due to the focus on ease of use and generalizability, `m2g` only requires the path of the BIDs [8] organized input directory, the output directory, and which of the pipelines is to be run (with other parameters able to be specified by the user). Through calling `m2g` with the simplest parameters:

~~~
          m2g --<type> <input_directory> <output_directory>
~~~

where `<type>` is either `dwi` for the diffusion pipeline, `func` for the functional pipeline, or `both` for when both pipelines should be run, `m2g` is able to estimate connectomes. We used the `m2g` pipeline to successfully estimate all of the connectomes used in this manuscript with data consisting of both functional and diffusion MRI nifti files of varying voxel resolutions and acquisition methods (see `DatasetInfo.pdf` in the Supporting Materials for further details). `m2g` is also able to process scans with non-isotropic voxels without issue due to the reslicing performed in both the `m2g-d` and `m2g-f` pipelines utilizing Dipy’s reslice function with tri-linear interpolation [9]. As reslicing an image to a different resolution results in the creation of new voxels with different intensity values, it may cause concern that there is significant change to the information contained within the images and by extension the connectomes. However, it has been shown that such processing does not significantly change the information for both diffusion and functional MRI [10, 11].

In addition to visual inspection of quality assurance (QA) figures generated after each step in the pipeline, we used the discriminability metric, which evaluates the fraction of measurements from the same individual that are closer to one another than they are to the measurement of any other individual [12], as a benchmark to the performance of `m2g` during development. The `m2g-d` pipeline was optimized on the Kirby21 dataset [13], while the `m2g-f` pipeline was optimized on both the Kirby21 and IBATRT [14] datasets because they are commonly used for test-retest analysis. Both pipelines were then validated using data from the Consortium of Reliability and Reproducibility (CoRR), consisting of 35 different studies from nearly 20 different institutions around the world, spanning the Americas, Europe, and Asia [15]. The CoRR data collection efforts were not harmonized, and all data (regardless of quality) were requested to be shared. This data repository was thus well-suited to test the robustness of our pipeline. Several additional open-access datasets from other repositories with different acquisition details, such as ABIDE [16, 17], were also processed with `m2g`.

The neuroparc repository of atlases, which we cultivated alongside `m2g`, was used as the main resource for the standardized parcellations used in `m2g`’s development [5]. This repository served to consolidate atlases from a multitude of sources, each registered to MNI152 space at 1, 2, and 4 mm^3^ voxel sizes [18]. Of the atlases listed there, 35 were used during the development of `m2g`. While neuroparc was developed in unison with `m2g`, the `m2g` pipeline is capable of utilizing any parcellation atlases provided by the user, provided it has been registered into MNI152 space [18].

Due to the lack of gold standards to determine the accuracy of an estimated connectome, certain forms of validation of the `m2g` pipeline are impossible. However, we have used surrogate metrics for QA to increase confidence in the outputs. For each MRI, `m2g` ran to completion while passing a basic QA metric consisting of (1) each connectome is a single connected component, (2) for all applicable datasets, discriminability values are above 0.7, (3) QA figures were generated successfully, (4) homophyly and homotopy observations are in agreement with literature. `m2g`’s ability to pass these metrics for validation shows its ability to be used widely, across a variety of datasets, including those leveraging older MRI technology.

To visually inspect consistency in estimated connectomes across the datasets, a distance-dependent group consensus structural connectivity graph was created. This method serves to prevent the observed trend of down-weighting long-range connections [19] in conventional averaging of connectomes and preserves the network properties seen in individuals. A similar graph was created for the functional MRI data, called a group-averaged functional connectivity matrix [20], which serves to minimize the effect of inconsistent correlations (Figure 2). The reproducibility of the estimated connectomes was tested using discriminability [12], and network statistics to determine that biological diversity, i.e. structural or functional properties unique to each subjects, and connectome plausibility were maintained.

**Figure 1.**
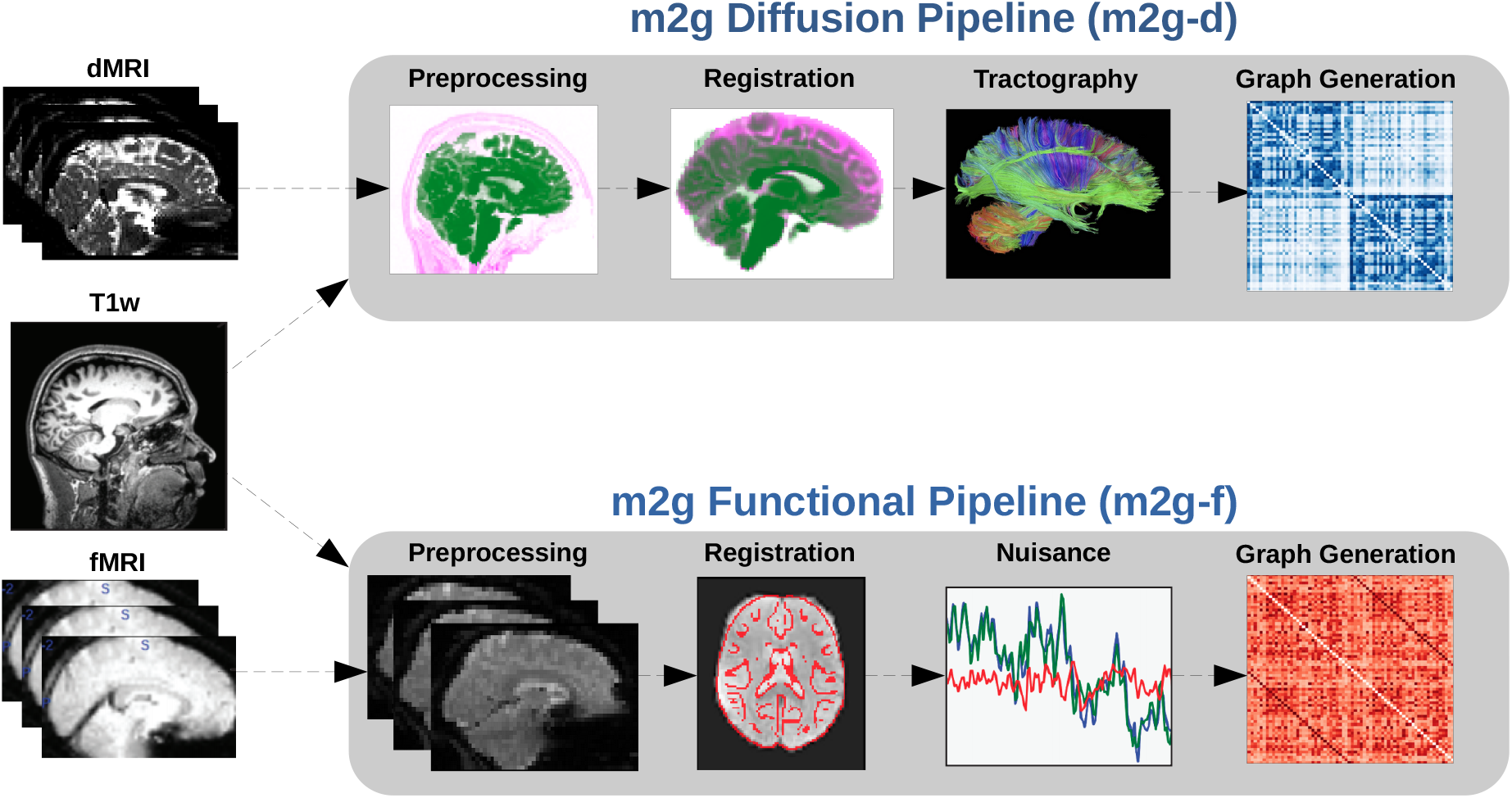
Individual Level Pipeline. The `m2g` pipeline has two sub-pipelines: `m2g-d` transforms Nifti-formatted dMRI data into sparse structural connectomes, and `m2g-f` organizes the data for processing by CPAC’s functional pipeline that we developed here. Each sub-pipeline consists of four key steps, and each step generates both data derivatives and quality assurance figures to enable both qualitative assessments and quantitative comparisons. Detailed descriptions of the processes involved in each step can be found in the m2g Diffusion Workflow and m2g Functional Workflow sections.

**Figure 2.**
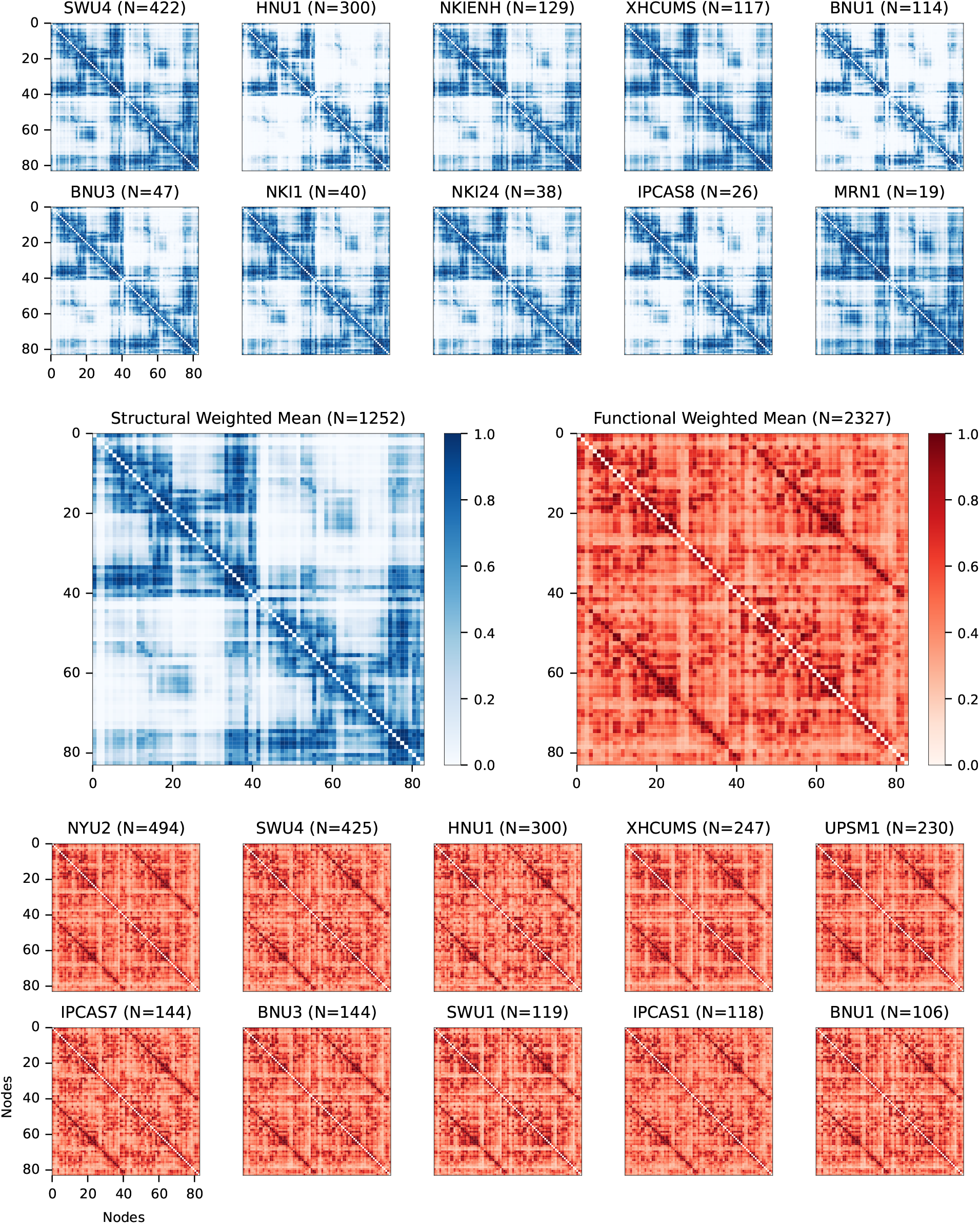
Group Consensus Connectomes. Group consensus structural connectome from `m2g-d` (blue) and group consensus functional connectomes from `m2g-f` (red), using the DKT parcellation method [21]. Structural connectomes of the datasets appear qualitatively similar, with minor deviations particularly visible in the contralateral regions of the connectomes (nodes 0-40 and 41-80). Ipsilateral connectivity is consistently more dense than contralateral connectivity in structural connectomes [22]. The functional connectomes appear qualitatively similar to one another. Homotopic correlation is consistently higher than ipsilateral and contralateral connectivity, which agrees with existing knowledge about functional correlation in the brain [23–25].

### 2.2 Validated Connectomes

Connectomes were estimated by both `m2g-d` and `m2g-f` for 35 parcellations from the Neuroparc repository [5] (see `Parcellations.pdf` in Supporting Materials). As mentioned before, the “true” accuracy of estimated connectomes compared to the true functional and structural properties of individuals’ brains is impossible to measure. Instead, we measure the reliability of the `m2g` pipelines through the discriminability of each dataset (Figure 3), and evaluate validity through various proxies, including the amount of homophyly and homotopy between regions of interest (Figure 4), and manual inspection of extensive QA images generated during each major step of the pipelines (see Appendix for example Figures 8-11). The connectomes estimated by `m2g` consistently, regardless of parcellation method/dataset, had corresponding discriminability metrics above random chance and in certain conditions achieving optimal results. Analysis of the homophyly and homotopy from connectomes estimated by `m2g` were found to agree with well documented phenomena [22–25], and manual inspection of a random sampling of QA images found no major errors.

**Figure 3.**
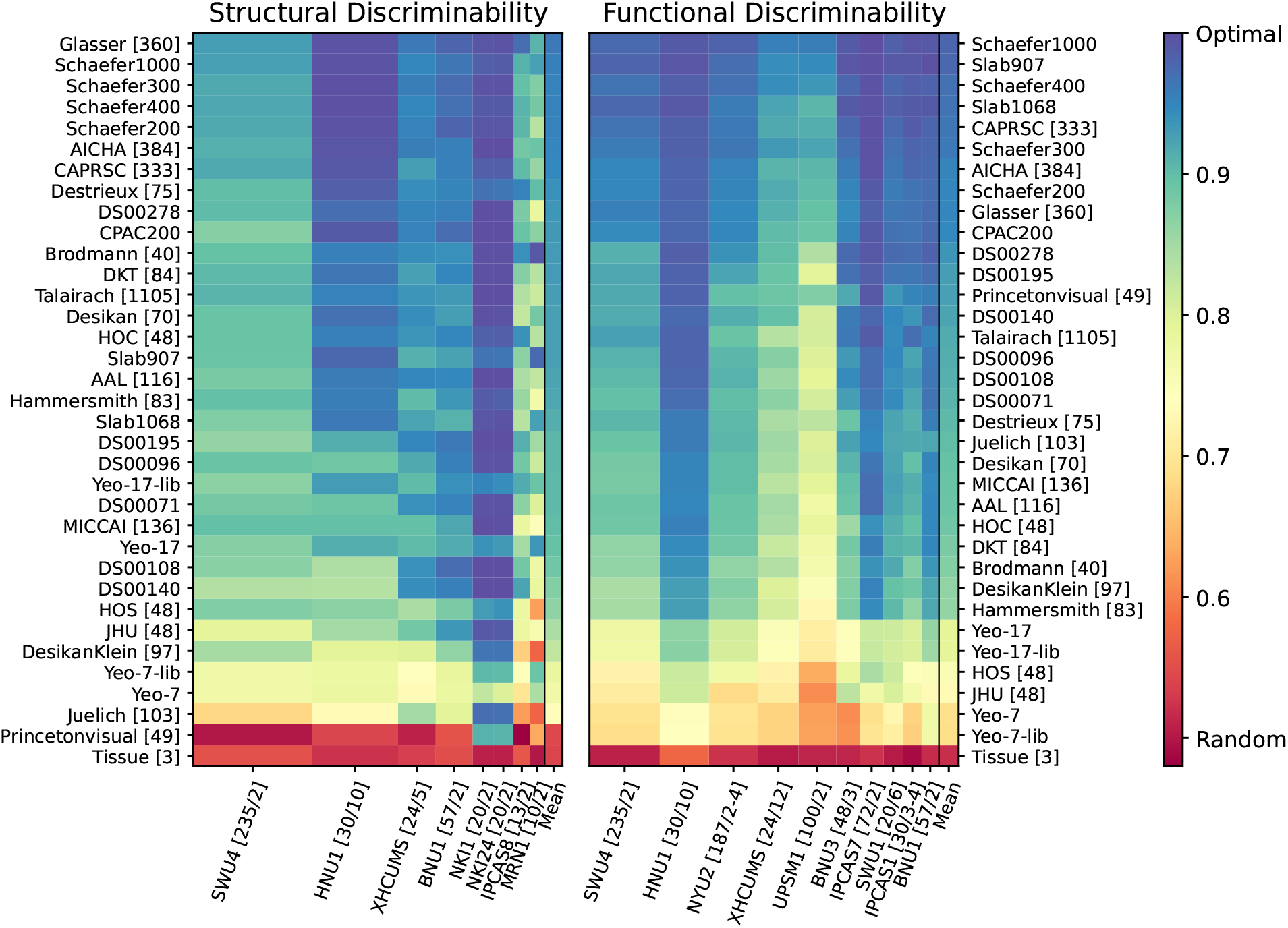
Discriminability Results. Discriminability values from each applicable diffusion (left) and functional (right) MRI dataset. Relative row height denotes the relative size of the dataset. Rows, each representing a different parcellation, are organized from top to bottom by highest to lowest mean discriminability value. The mean discriminability value for each parcellation is displayed in the last column of both plots. The number of subjects/sessions is displayed next to the datasets’ names in brackets and the number of ROI’s in a given parcellation are shown in brackets if not mentioned in the parcellation name. Discriminability values for the structural connectomes was greater than 0.7 for the 32 of the 35 parcellations, while being robust to the number of scans per subject. Functional connectomes rarely had discriminability values lower than 0.7.

**Figure 4.**
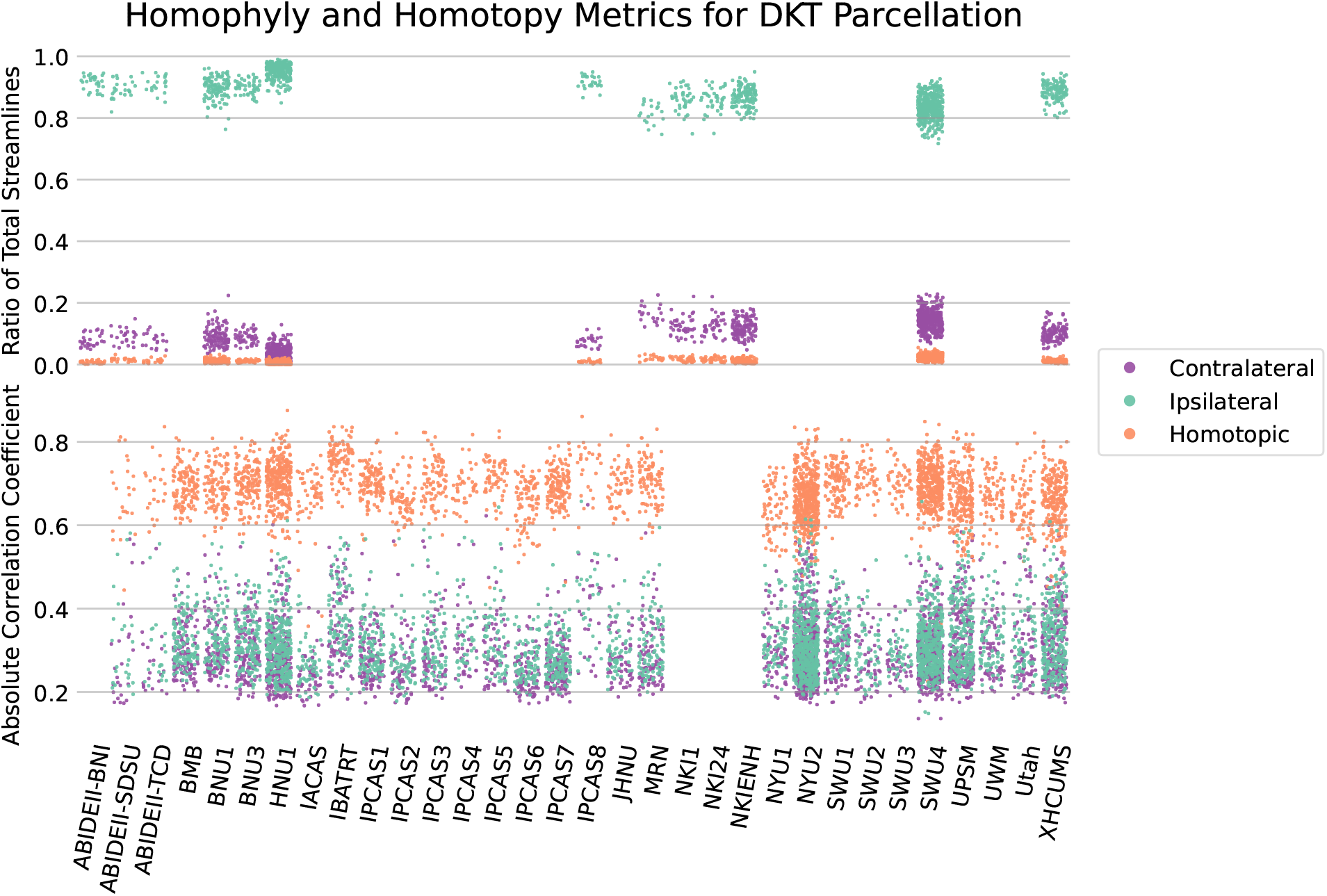
Connectome biological plausibility results. Analysis of the edge weights of both structural (top) and functional (bottom) connectomes estimated using the DKT parcellation method. For the structural connectomes, the mean of the percentage of streamlines observed between ipsilateral, contralateral, and homotopic ROIs was recorded and plotted for each scan. Mean Pearson correlation coefficients between ipsilateral, contralateral, and homotopic ROIs was plotted for functional connectomes. A consistent significantly higher ratio of ipsilateral connections was observed across parcellation methods for structural connectomes, as well as a higher correlation between homotopic ROIs across parcellation methods in functional connectomes.

#### Discriminability

Discriminability [12] was estimated for every dataset with multiple scans per subject (as reported in the Supporting Materials, which gives all of the individual numeric values). The closer to 1 the discriminability value of a processed dataset is, the more distinct the connectomes estimated from one individual’s scans are from the connectomes of others. A discriminability value of one supports the estimated connectome capturing some unique functional or structural property for each of the subjects. The `m2g-d` pipeline estimated connectomes with discriminability values greater than 0.7 for the 33 of the 35 parcellations (Figure 3, top). The discriminability measurements appear to be robust to the number of scans per subject, as can be seen with the SWU4 and HNU1 datasets (two scans per subject and ten scans per subject, respectively). In addition, `m2g-f` rarely estimated connectomes with discriminability values lower than 0.7 (Figure 3, bottom).

#### Connectome Connectivity

Because discriminability assesses the uniqueness of each individual’s connectome, it does not guarantee that the connectome itself is biologically meaningful. To address this issue, we analyzed the prominence of different categories of connections—ipsilateral versus contralateral and homotopic versus heterotopic—using the connectomes estimated from three unique parcellations. To analyze homotopic connections, we chose the DKT [21], AAL [26], and Hammersmith [27] parcellations, due to their symmetric ROI placement (i.e. each ROI on the left hemisphere had a matching ROI on the right). For the structural connectomes, the percentage of total edges which belonged to each network were analyzed, while the absolute Pearson’s correlation value [28] calculated by CPAC was used for the functional connectomes. The results coincided with well-documented phenomena regarding structural and functional relationships [22–25]. Namely, structural connectomes had significantly more ipsilateral than contralateral connections, while functional connectomes had significantly stronger homotropic correlations than heterotropic, either ipsilaterally or contralaterally (Figure 4).

#### Reproducibility

In the `m2g` pipeline, each processing procedure may introduce bias or noise in the estimated connectome. The randomized seed placement which takes place in the tractography step of the `m2g-d` pipeline, and the registration of the functional MRI data onto the T1-weighted image and the T1-weighted image onto the diffusion MRI data for `m2g-f` and `m2g-d`, respectively, serve as potential vectors for noise. To test the reproduciblity of `m2g`, we estimated connectomes for a subset of scans multiple times. Using a subset of five scans from each dataset, we used `m2g` to estimated connectomes five times per scan, using the same default parameters. The resulting connectomes were then treated as unique, with each set belonging to the same subject. Discriminability metrics were then calculated to determine whether the unique qualities of the individual scan were preserved. For all subsets, discriminability values of 1 were recorded for all parcellations for both the functional and structural connectomes. This supports the robustness of `m2g` to any bias or noise, as the unique properties of each subject are significant enough not to be altered. Additional testing using Spearman’s rank correlation found that the repeated connectomes had a coefficient greater than 0.98 for all parcellations.

#### Quality Assurance Outputs

Quality assurance images were created by `m2g` at each of the significant steps in both the `m2g-d` and `m2g-f` pipelines (Figure 1) for easy determination of erroneous results. The QA figure formats were chosen for clarity and minimal image analysis expertise requirements in order to detect any potential errors. A random sample of the QA images from 15 estimated connectomes per dataset were visually inspected and found no significant issues. Samples of the output QA images can be found in the Appendix. Documentation for each of the QA figures generated by `m2g` can be found at https://neurodata.io/mri/.

### 2.3 Low Resource and Time Requirement

Computational expediency and resource efficiency were two key reasons we developed `m2g`. While the required resources for running `m2g` varies according to the data, the datasets mentioned in this paper could be processed by `m2g` with approximately 16 GB of RAM and 1 CPU core. If more resources are available, `m2g` is capable of utilizing multiple CPUs at the same at various points throughout the pipeline, such as registration and graph generation, to reduce runtime. While estimating the connectomes used in this manuscript, we recorded the time required for `m2g` to successfully analyze each of the scans used in the discriminability calculations (Figure 5). The optimal configuration of resources for our purposes required 3 CPUs and 16 GB of RAM. When tested using 1 CPU, `m2g-d` took less than 120 minutes to complete the connectome creation using 35 parcellations for a given dMRI scan. The `m2g-f` pipeline likewise took less than 110 minutes to generate the set of connectomes. Additional CPUs were found to significantly increase the RAM requirements of the `m2g-f` pipeline, due to the configuration of CPAC.

**Figure 5.**
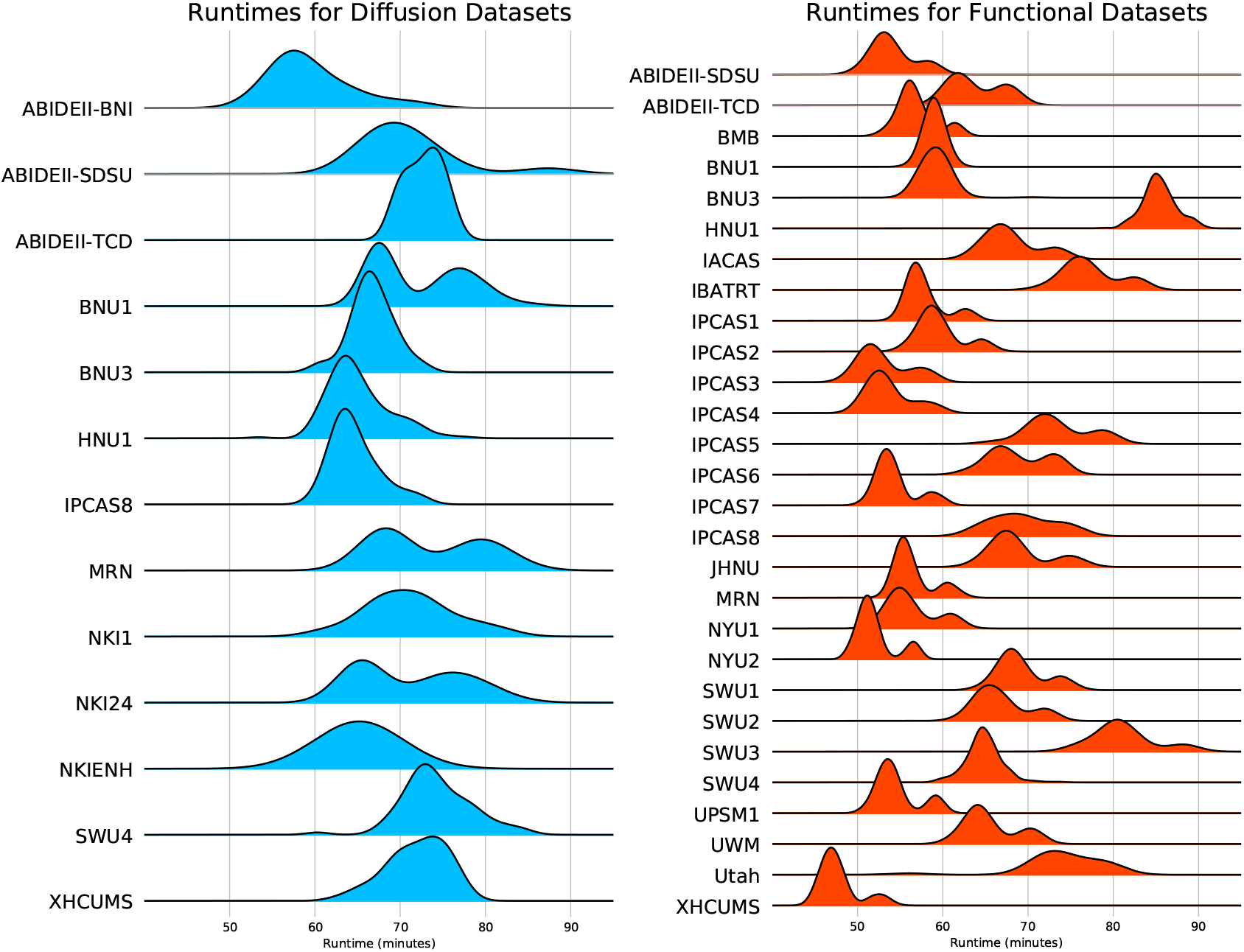
Runtimes for Diffusion and Functional datasets. Amount of time, in minutes, it took for `m2g` to generate a connectome for each of the scans in the datasets using the 35 brain parcellations. Computation time for diffusion datasets (blue) and functional datasets (red) varied based off of size and resolution of the input MRI files. In these time estimates, `m2g` was run utilizing 3 CPUs in parallel and at least 16 GB of RAM available.

## 3 Discussion

The `m2g` processing pipeline was created to both efficiently analyze diffusion and functional MRI scans and lower the barrier for entry to connectomics. It was developed to excel at three key aspects: ease of use, biological veracity, and computational reproducibility. We believe `m2g` to be a success with respect to these three aspects.

Attesting to its ease of use, all datasets referenced in this manuscript were run through `m2g` with the default parameters and minimal user input, resulting in the successful generation of connectomes for each scan. This was true across a range of acquisition properties of the MR images and with modest computational resources: less than two hours for each scan using 16 GB of RAM and 3 CPUs and no intra-pipeline additional parallel processing steps. In addition, `m2g` requires very minimal user input: only the input/output locations, pipeline type, and acquisition method (for fMRI data). As such, the pipeline is an easy tool for researchers and clinicians without extensive computer science experience or resources. Also, `m2g` produces comprehensive visual reports to analyze processing output: we designed the QA figures to easily visually determine whether the pipeline has successfully processed the provided data. With the emphasis on color coordination and concise figure generation, many issues in the preprocessing and registration process is easily visible. All QA figures are clearly labeled and placed in the same directory location relative to the output, regardless of pipeline being used, with detailed explanations found at http://m2g.io.

We assessed `m2g`’s ability to produce biologically plausible and realistic data using multiple approaches. First, and most simply, we determined that there was a path from each ROI to each other ROI, i.e., that the brain was connected, which we know to be generally true. Second, our visual inspection of output QA figures showed no obvious abnormalities that would negatively effect the accuracy of the connectome. Third, we computed the discriminability for each dataset. Discriminability surpassed statistical tests of random chance for essentially all parcellations across datasets. In fact, the majority of the 35 parcellation methods used in this analysis resulted in discriminability values above 0.8 (Figure 3). Fourth, we assessed the relative fraction of ipsilateral versus contralateral, and homotopic versus heterotopic connections. Results were consistent with current knowledge about the anatomical and functional realities of neuro-typical brains [22–25]. More specifically, the prominence of ipsilateral connections in structural connectomes, and the prominence of homotopic correlations in functional connectomes (Figure 4).

Our focus on computational reproducibility manifested through `m2g` being open source and fully supporting execution via the Docker container platform. From version to version of `m2g`, prospective users do not need to worry about reproducibility issues due to dependency issues as container images are generated and uploaded to a public repository for each new version of m2g. These containers are released with version-controlled libraries for m2g and all its dependencies, maximizing run-to-run reproducibility in an easy way. The Docker image for m2g can be found at https://neurodata.io/mri/. We also confirmed within version computational reproducibility for `m2g` through the re-estimation of connectomes, which showed optimal discriminability values and Spearman’s rank correlation greater than 0.98 for essentially all parcellations across datasets.

The design criteria for `m2g` required certain trade-offs in performance to increase the generalizability of input data. With the strength of pipeline flexibility and minimal required user input still being paramount, `m2g` could be improved along several dimensions. First, recent advances in registration [29] and tractography [30] could be incorporated. Second, several more sophisticated batch effect strategies have been successfully employed in dMRI data [31, 32]. Such strategies could possibly help here as well, especially if they are modified appropriately to work on binary graphs [33]. Third, `m2g` may under-perform for particular populations (e.g., infants), for brains that show nonstandard structures such as tumors, resected regions, lesions, or other species (e.g., non-human primates [34]). Inclusion of the ability to supplement the standard MNI152 reference images and parcellations with ones relevant to nonstandard brain structures could drastically expand the populations `m2g` could process. This would be particularly interesting as a means to adapt the workflow to data collected from rodents and nonhuman primates in the future.

Because the methods developed during the creation of `m2g` are open-source and easy to use, and the data are open-access, the continual development of the pipeline and assessment of the connectomes generated is ready for outside collaboration. Due to the well-commented and modular code, modification is straightforward for python users. Along with the ease of use, a boon for the wider adoption of `m2g` is the plethora of connectomes created as a byproduct of its development. These connectomes are open-access and available from our website, https://neurodata.io/mri/.

## 4 Methods

### 4.1 Data

The `m2g` pipelines were validated through the processing of the majority of data from the Consortium of Reliability and Reproducibility (CoRR). The CoRR data consists of 36 different studies from nearly 20 different institutions around the world, spanning the Americas, Europe, and Asia [15]. The CoRR data collection efforts were not harmonized, and all data (regardless of quality) were shared. Thus, these collections are well-suited to test the robustness of any pipeline. In addition to the CoRR data, we also used `m2g` to process several additional open-access data collections with complementary acquisition parameters (see Supporting Material `DatasetInfo.pdf`). Using this collection of MRI data, `m2g` pipeline development and parameter selection prioritized the maximization of discriminability scores across all datasets, while also focusing on limiting resource requirements.

### 4.2 m2g Input

The input required to run the `m2g` pipeline depends on whether functional and/or diffusion MRI data is being analyzed. The required inputs for either pipeline are:

- A T1-weighted anatomical scan, either an uncompressed or gzipped nifti file
- `m2g-d`:
  - A diffusion-weighted MRI file, either an uncompressed or gzipped nifti file
  - b-value and b-vector parameter files (.bval and .bvec file types)
- `m2g-f`:
  - A functional MRI file, either an uncompressed or gzipped nifti file
  - The TR value (in seconds) and acquisition method used for the functional scan

The directory containing the input data must be BIDS formatted [8]. Both the `m2g-d` and `m2g-f` pipelines can be run independently or on the same dataset, provided the required data files are available. Which parcellation(s) to be used in the connectome creation can be specified by the user, whether they be unique files stored locally or from the neuroparc repository [5]. If no parcellations are present, then `m2g` will pull from the neuroparc repository and run all available parcellations.

### 4.3 m2g Diffusion Workflow

The `m2g` diffusion pipeline leverages existing open source tools, including the fMRI Software Library (FSL) [35–37], Dipy [9], the MNI152 atlas [18], and a variety of parcellations defined in the MNI152 space [5]. The pipeline consists of five major steps: (1) Preprocessing, (2) Registration, (3) Tensor estimation, (4) Tractography, and (5) Graph generation. All algorithms used throughout the pipeline requiring hyper-parameter selection were initially set to the suggested parameters, and optimized on the subset of CoRR datasets. The output of each processing stage includes data derivatives and QA figures to enable individualized accuracy assessments. The QA figures at each major step include cross-sectional images at different depths in the three canonical planes (sagittal, coronal, and axial) of images or overlays. Figure 1 provides a schematic of the individual-level analysis.

#### Preprocessing

The input diffusion MRI data is first eddy-corrected using FSL’s eddy_correct [9] program with the default parameters. This program was chosen as newer eddy functions either require substantially longer to run or rely on GPU acceleration, an additional resource requirement which would reduce the accessibility of `m2g`. The associated bvec and bval files are then checked for any errors and, if necessary, corrected using Dipy’s read_bvals_bvecs function [9] into a usable format. The b-vectors are then normalized, making sure to not alter the b-vector if its associated b-value is 0.

The eddy-corrected dMRI file then has its orientation checked, and if not already oriented in RAS+ coordinate space, nibabel’s as_closest_canonical [38] is used for the reorientation. The b-vectors are also reoriented to RAS+ if necessary using default parameters. The dMRI file is then resliced to the desired voxel size (specified by the user, with a reasonable default of 2mm^3^ if not specified) using Dipy’s reslice function with trilinear interpolation[9].

The T1 weighted MRI file (T1w) is also reoriented into RAS+ format using as_closest_canonical and similarly resliced, if necessary, into the desired voxel size using Dipy’s reslice function [9] with default parameters. The resulting image is then skull-stripped using AFNI’s 3dSkullStrip function [39] with the surface density parameter set to 30, resulting in an anatomical image with all non-neuronal structures removed.

#### Registration

From the skull-stripped T1w anatomical file, three probability masks are generated for the grey matter, white matter, and cerebral spinal fluid regions of the brain using FSL’s fast function [35–37] using default parameters. The white matter mask is then used to create another mask of just the outer edge of the white matter area using FSL’s fslmaths [35–37]. Using FSL’s flirt function [35–37], an initial linear registration is made for the anatomical image to the reference MNI152 template in MNI space. An initial attempt is made for non-linear registration of the T1w image to the MNI template, using the affine transformation matrix previously created by the linear registration as a starting configuration. If it is not successful and throws an error, linear registration is used. Using FSL’s fnirt function [35–37], the nonlinear warp coefficients/field is generated to register the T1w image onto the reference MNI image.

FSL’s invwarp function [35–37] is then used on the coefficients/field to get the field for registering the reference MNI image onto the T1w image. However, if the nonlinear registration fails, then the linear registration affine transformation matrix for T1w to MNI is used and inverted using FSL’s convert_xfm [35–37].

FSL’s flirt is then used for calculating the boundary-based registration (BBR) of the dwi image onto the T1w image, using 256 bins and 7 degrees of freedom. The resulting transformation matrix is inverted and used with a mutual information cost function in flirt to generate an image of the anatomical image registered to the dwi image. Using fslmaths, binary ventricle and corpus callosum masks in MNI space are generated. Along with the previously generated grey matter, white matter, and cerebral spinal fluid masks, flirt is used to register the masks to dwi space. A grey matter/white matter boundary binary mask is then made using fslmaths, which denotes the approximated boundary between the two tissue types across the entire brain, for later seed propagation in the tractography stage of the pipeline.

Finally, the parcellation files to be used in the graph generation are reoriented using nibabel’s as_closest_canonical function into RAS+, where they are resliced to the desired voxel size using Dipy’s reslice function. They are then registered from MNI to dwi space using the previously calculated transformation matrices, utilizing FSL’s flirt function and the “nearest neighbor” cost function.

#### Tractography

Seeds are generated along the grey matter/white matter boundary mask of the registered brain using a custom python script, where the number of seeds per voxel on the mask can be specified by the user. The seed locations and dwi image are then used in conjunction with Dipy tractography functions [9] in order to generate a series of non-directional streamlines spanning the image. `m2g` offers deterministic or probabilistic tractography, Constrained Spherical Deconvolution (CSD) or Constant Solid Angle (CSA) reconstruction methods, and particle filter or local tracking. Regardless of the tractography settings specified, the resulting series of streamlines is output and saved to a trk file for use in graph generation.

#### Graph Generation

For each parcellation being analyzed, each streamline is converted into voxel coordinates and analyzed for overlap with any of the regions specified by the parcellation. `m2g` does not determine the direction of a given streamline, making its connectome and resulting graph undirected. If a streamline overlaps with a given region for more than 2mm, it is considered an intersection. The number of streamlines intersecting a given region and the other regions they also intersect with are recorded and tabulated in the form of a weighted edgelist. In this edgelist format each unique region of interest on the parcellation, marked by a unique intensity value, is considered a node whose edge weight is the number of streamlines that connect it to another region. For example, if a streamline intersects two or more regions in the parcellation (region A, region B, and region C) the weight of the edges AB, AC, and BC is increased by one. In this sense, the order through which the streamline intersects is not important. The resulting edgelist is saved in the form of a csv file, and an accompanying adjacency matrix of the normalized edgelist is also saved. A unique edgelist will be made for each parcellation specified by the user.

### 4.4 m2g Functional Workflow

The `m2g-f` pipeline was constructed starting with the optimal processing pipeline identified in Wang et. al [12] using CPAC [6]. The CPAC pipeline utilizes existing open source tools, including FSL [35–37], the MNI152 atlas [18], and a variety of parcellations defined in the MNI152 space [5]. The functional pipeline consists of four major steps: (1) Preprocessing, (2) Registration, (3) Nuisance correction, and (4) Graph generation.

#### Preprocessing

The `m2g-f` pipeline in CPAC uses AFNI’s SkullStripping function [39] with a variable shrink factor between 0.4 and 0.6 over 250 iterations with nearest neighbor interpolation to eliminate all non-neuronal structures from the anatomical image. The resulting anatomical file is resampled to the desired resolution voxel size using FSL’s FNIRT [35–37]. Slice timing correction is then performed using AFNI’s 3dTshift [39], utilizing the TR value and scan acquisition method provided by the user. The resulting image is then motion corrected using AFNI’s 3dvolreg, aligning all images to the first image in the functional MRI’s timeseries.

#### Registration

Nonlinear boundary based registration of the preprocessed fMRI and anatomical images are performed in order to transform them into MNI152 space. The MNI152 6th generation anatomical reference image [18] is used in this registration process, as it is FSL’s preferred image. The functional image is then registered to the anatomical scan using FSL bet [35–37] with BBR. FSL’s standard white matter, grey matter, and cerebral spinal fluid masks are then registered onto the functional image using FSL’s FAST thresholding, resulting in segmentation tissue masks. The specified atlases that are going to be used in the connectome generation are also registered to the functional space using FSL’s FLIRT function.

#### Nuisance Correction

Nuisance correction of the functional image is performed using the component based noise correction (CompCor) method. Using “noise ROIs”, or areas of white matter and cerebral spinal fluid whose intensity values are unlikely to be modulated by neural activity, physiological noise can be isolated[40].Based on this assumption, physiological noise in gray matter regions can be corrected for by regressing out principal components from noise ROIs. CompCor uses the previously registered white matter and cerebral spinal fluid masks (csf) in order to determine noise ROIs. A principal component analysis is applied using the top five components of the white matter and csf to characterize the times series data from the noise ROIs. Significant principal components are then introduced as covariates in a general linear model as an estimate for the physiological noise single space. Polynomial Detrending is also performed to remove linear or quadratic trends in the timeseries, likely from changes in scanner heat or subject movement. After nuisance regression, frequency filtering occurs on the functional data using a bandpass filter from 0.01 Hz to 0.1 Hz to account for low-frequency drift and high-frequency noise.

#### Graph Generation

With the anatomical files and atlases registered to the functional image, the estimation of the functional connectomes for each atlas can occur. For each ROI, the timeseries of the average of all voxels within the ROI at each collection time point is calculated. This timeseries is then used to calculate the Pearson’s correlation coefficient [28] between each pair of ROIs. The functional connectome is then rank-transformed by replacing the magnitude of the correlation with its relative rank, from smallest to largest [12]. The resulting adjacency matrix of correlations is then saved, along with an edgelist file containing the same information.

### 4.5 Validation of m2g on Diverse Data

#### Discriminability

To evaluate a method’s reliability, Wang et al. [12] developed a metric called “discriminability” that quantifies the fraction of measurements from the same individual that are closer to one another than they are to the measurement of any other individual. Discriminability, as seen in Equation (4.1), describes the probability that two observations within the same class are more similar to one another than two observations belonging to a different class:

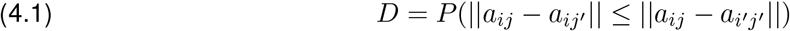

In the context of validation of `m2g`, each connectome is converted into a ranked vector, assigning each edge a weight between 0 and 1 from smallest to largest, and compared. This means that each connectome *a*_*ij*_, in a test-retest dataset is first compared to other connectomes belonging to the same subject, *a*_*ij*_*′*, and then to all connectomes belonging to other subjects, *a*_*i*_*′*_*j*_ *′*. A perfect discriminability score is 1, meaning that for all observations within the dataset, each connectome is more alike to connectomes from the same subject than to others. Optimizing `m2g` with respect to discriminability enables us to minimize the upper-bound on error for any general downstream inference task. The discriminability score for the multitude of datasets run through the default settings of `m2g` were recorded (see Supporting Material `DiscrimNumbers.xlsx`). The average discriminability over all scans was approximately 0.85 for dMRI data and approximately 0.8 for fMRI data.

#### Connectome Connectivity

While discriminability may determine the extent that `m2g` preserves the unique properties of a given MRI scan, additional care must be taken to determine whether the connectomes generated by `m2g` conform with basic neurological phenomena. One way to do this involves observation of either the physical or functional connections between different regions of the brain. Three simple categories for these regions are ipsilateral (connections that stay within the left or right hemispheres), contralateral (across hemispheres), or homotopic (from one region of the brain to the same region on the other hemisphere). Using the edge weights from the structural connectomes generated by `m2g`, the percentage of edges that fell into each of the three connection types was calculated, and the mean and standard deviation were found for each dataset. This process was performed for three parcellations which had clearly-defined homotopic regions of interest, namely the DKT, Hammersmith, and AAL atlases. For the functional connectomes generated, the Pearson’s correlation between regions of interest was used as a metric. The mean and standard deviation of each of the three connection types was found for each dataset (Figure 4).

## 5 Ethical Compliance

We complied with all relevant ethical regulations. This study reused publicly available data acquired at many different institutions. Protocols for all of the original studies were approved by the corresponding ethical boards.

## 6 Software Availability

All of our processed data is available from our website, http://m2g.io, and has been deposited into our public github repository, https://github.com/neurodata/m2g, and published with a DOI, https://doi.org/10.5281/zenodo.1161284, under the Polyform License. Detailed documentation can be found at https://docs.neurodata.io/m2g/.

## 7 Data Availability

The data derivatives that support the findings of this study are available from our website, http://m2g.io, under a (ODC-By) v1.0 license.

## Acknowledgements

The authors from JHU are grateful for the support by the XDATA program of the Defense Advanced Research Projects Agency (DARPA) administered through Air Force Research Laboratory contract FA8750-12-2-0303; DARPA SIMPLEX program through SPAWAR contract N66001-15-C-4041; DARPA GRAPHS contract N66001-14-1-4028; National Science Foundation grant 1649880, and the Kavli Foundation for their support. Dr. Xi-Nian Zuo received funding support in China from the National Basic Research (973) Program (2015CB351702), the National R&D Infrastructure and Facility Development Program “Fundamental Science Data Sharing Platform” (DKA2017-12-02-21), the Natural Science Foundation of China (81471740, 81220108014) and Beijing Municipal Science and Tech Commission (Z161100002616023, Z161100000216152). Dr. Calhoun received funding from the NIH (P20GM103472 and R01EB020407) and the NSF (grant 1539067).

## 8 Appendix

**Figure 6.**
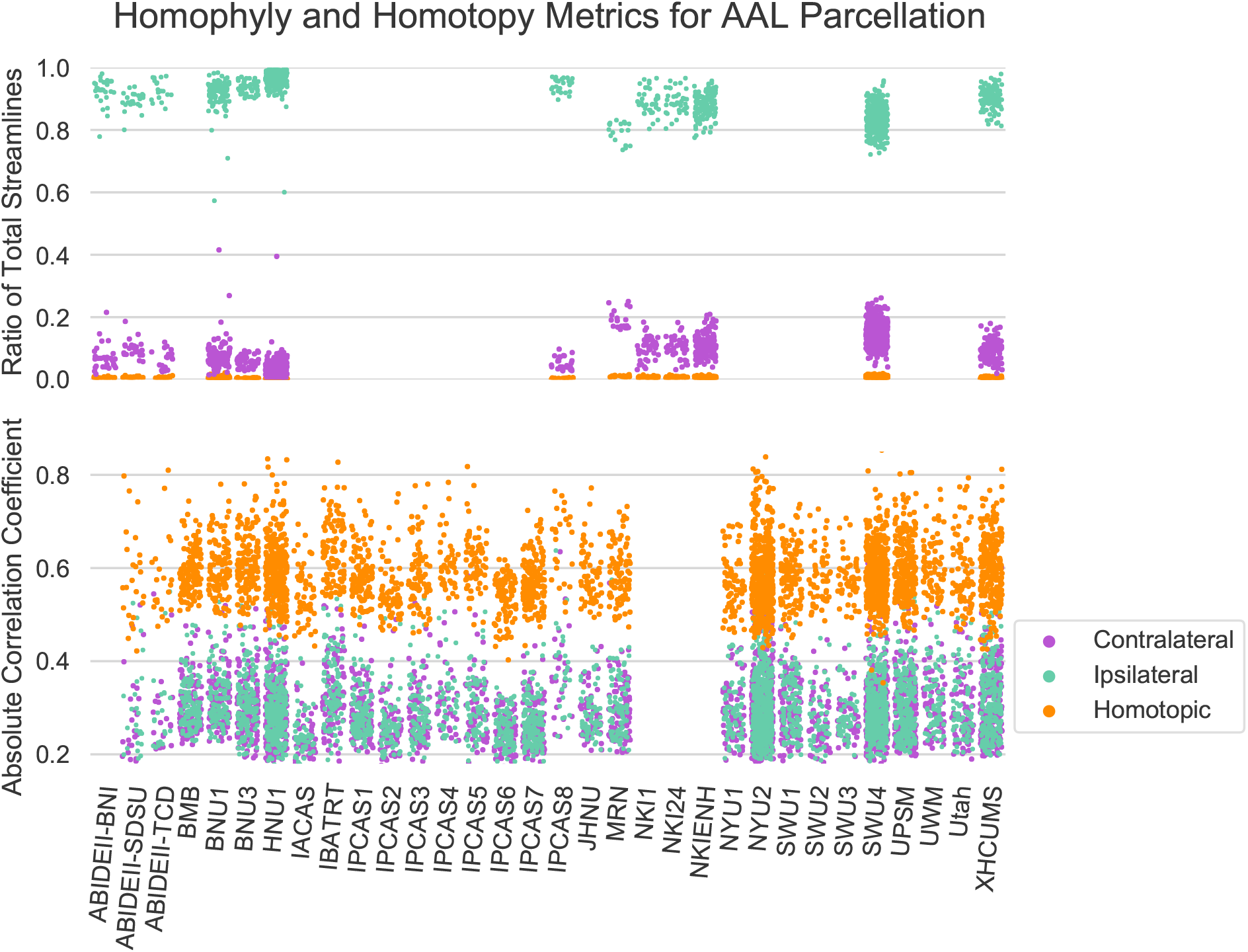
Connectome biological plausibility results. Analysis of the edge weights of both structural (top) and functional (bottom) connectomes estimated using the AAL parcellation method. For the structural connectomes, the mean of the percentage of streamlines observed between ipsilateral, contralateral, and homotopic ROIs was recorded and plotted for each scan. Mean Pearson correlation coefficients between ipsilateral, contralateral, and homotopic ROIs was plotted for functional connectomes. A consistent significantly higher ratio of ipsilateral connections was observed across parcellation methods for structural connectomes, as well as a higher correlation between homotopic ROIs across parcellation methods in functional connectomes.

**Figure 7.**
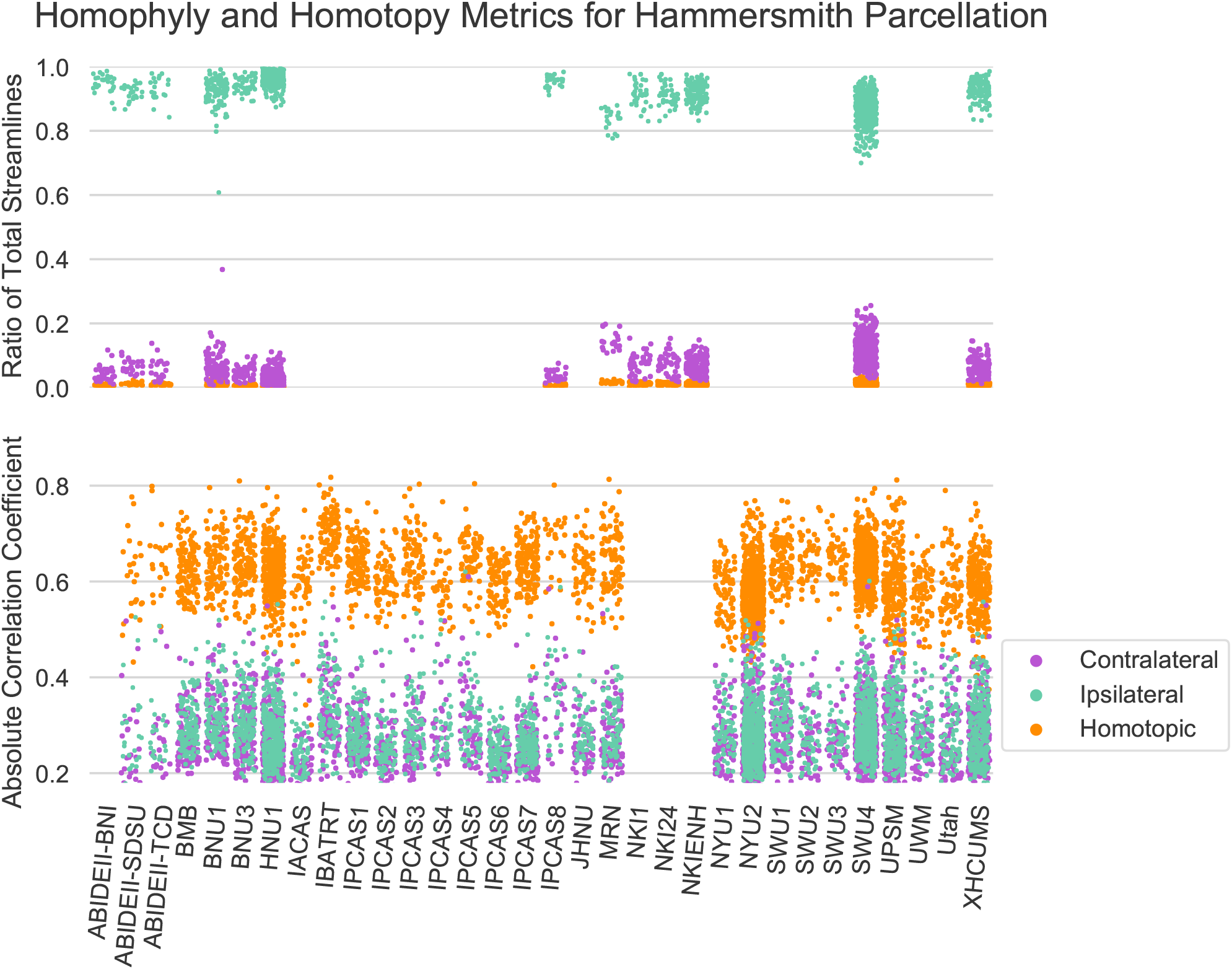
Connectome biological plausibility results. Analysis of the edge weights of both structural (top) and functional (bottom) connectomes estimated using the Hammersmith parcellation method. For the structural connectomes, the mean of the percentage of streamlines observed between ipsilateral, contralateral, and homotopic ROIs was recorded and plotted for each scan. Mean Pearson correlation coefficients between ipsilateral, contralateral, and homotopic ROIs was plotted for functional connectomes. A consistent significantly higher ratio of ipsilateral connections was observed across parcellation methods for structural connectomes, as well as a higher correlation between homotopic ROIs across parcellation methods in functional connectomes.

**Figure 8.**
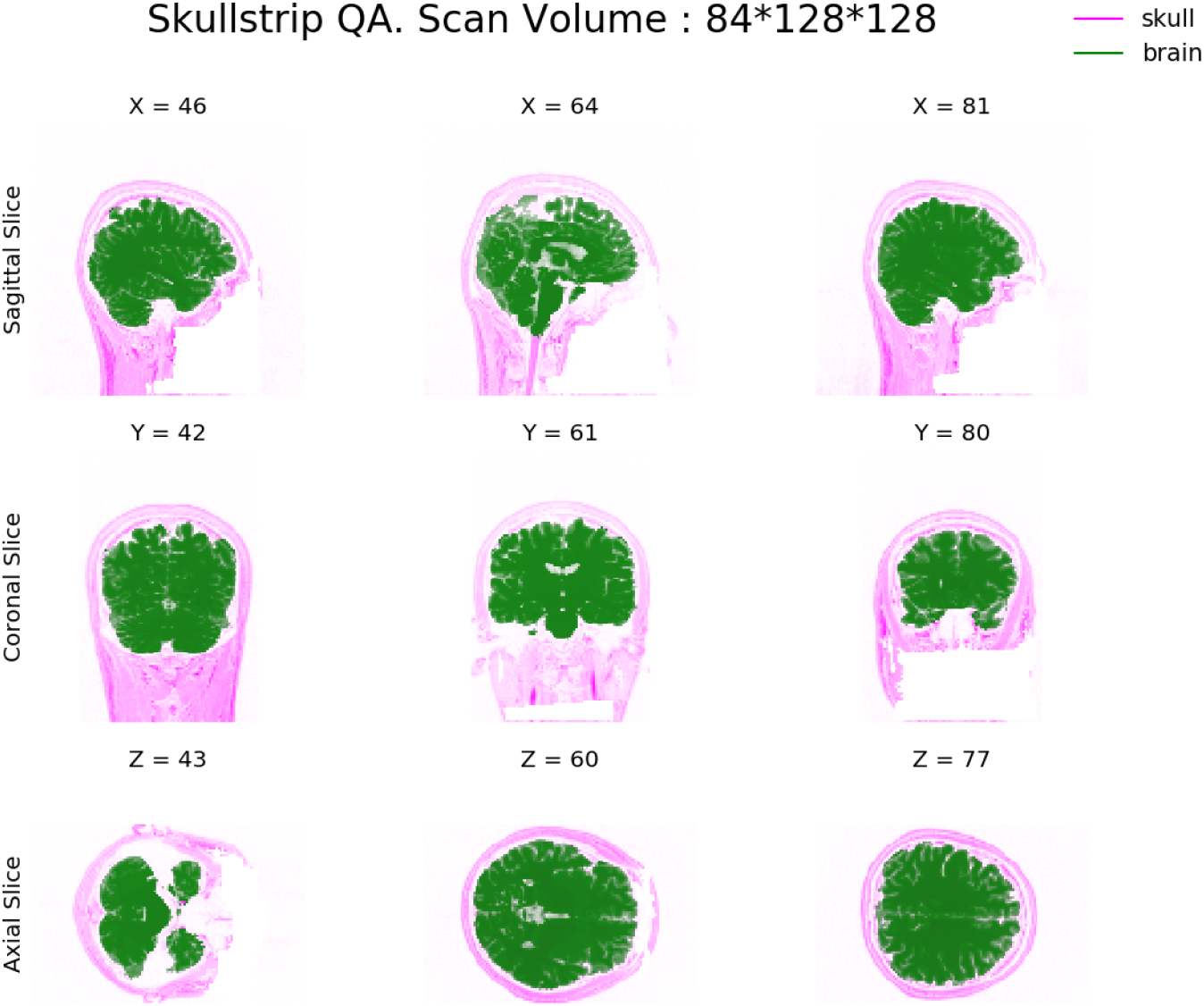
Skull-stripping. `m2g-d` **QA** Quality assurance figure from the `m2g-d` pipeline displaying the skull-stripped brain (green) over the original anatomical image (magenta). Images are sampled from three locations along the sagittal, coronal, and axial reference directions.

**Figure 9.**
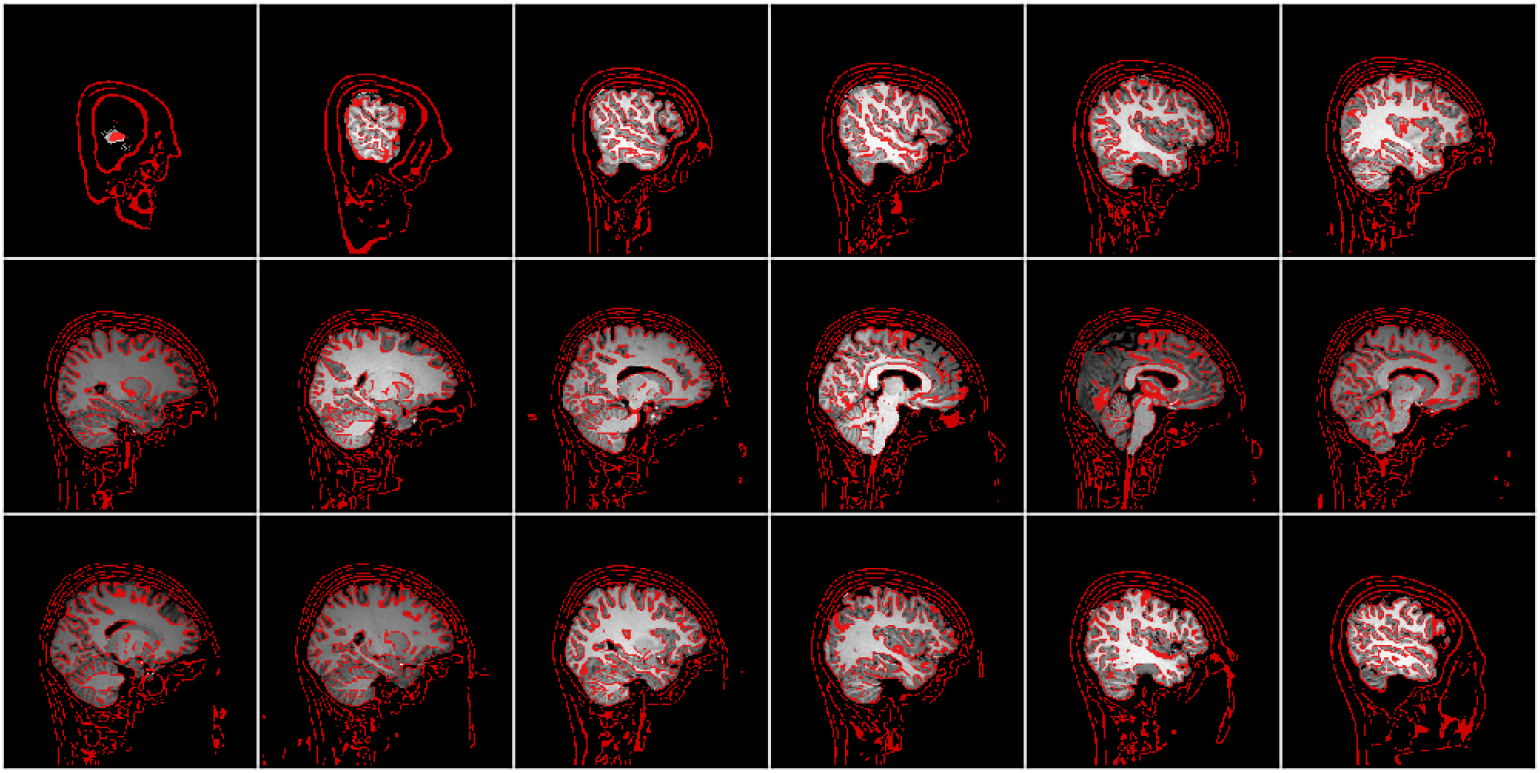
Skull-stripping. `m2g-f` **QA** Quality assurance figure from the `m2g-f` pipeline displaying the skull stripped brain (white) and the outline of the removed skull (red). This particular output displays images from the sagittal reference direction, with another output displaying from the axial direction.

**Figure 10.**
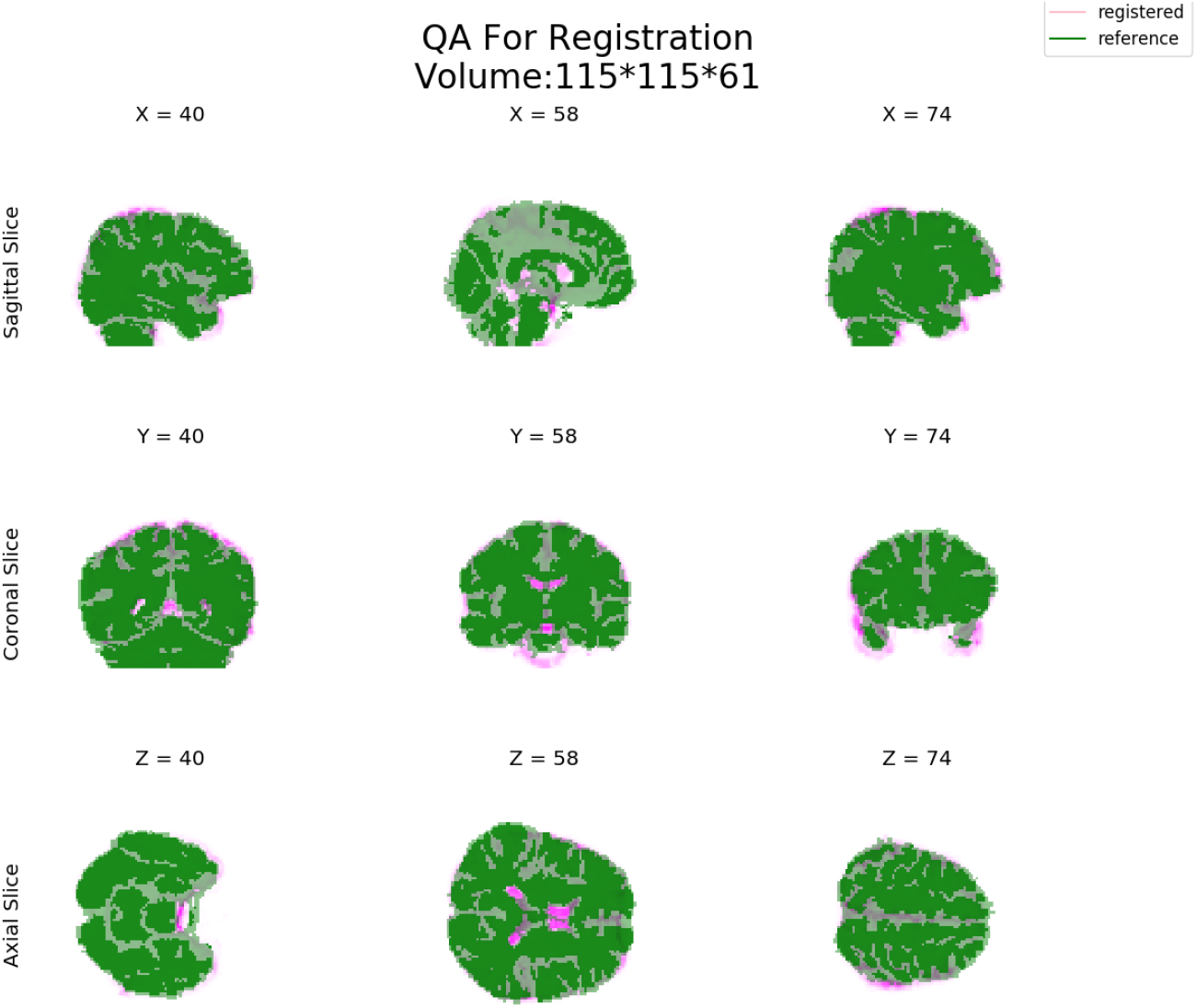
Parcellation Registration QA. Quality assurance figure from the `m2g-d` pipeline displaying the registered Tissue parcellation (green) over the brain it is registered to from the anatomical image(magenta). Images are sampled from three locations along the sagittal, coronal, and axial reference directions.

**Figure 11.**
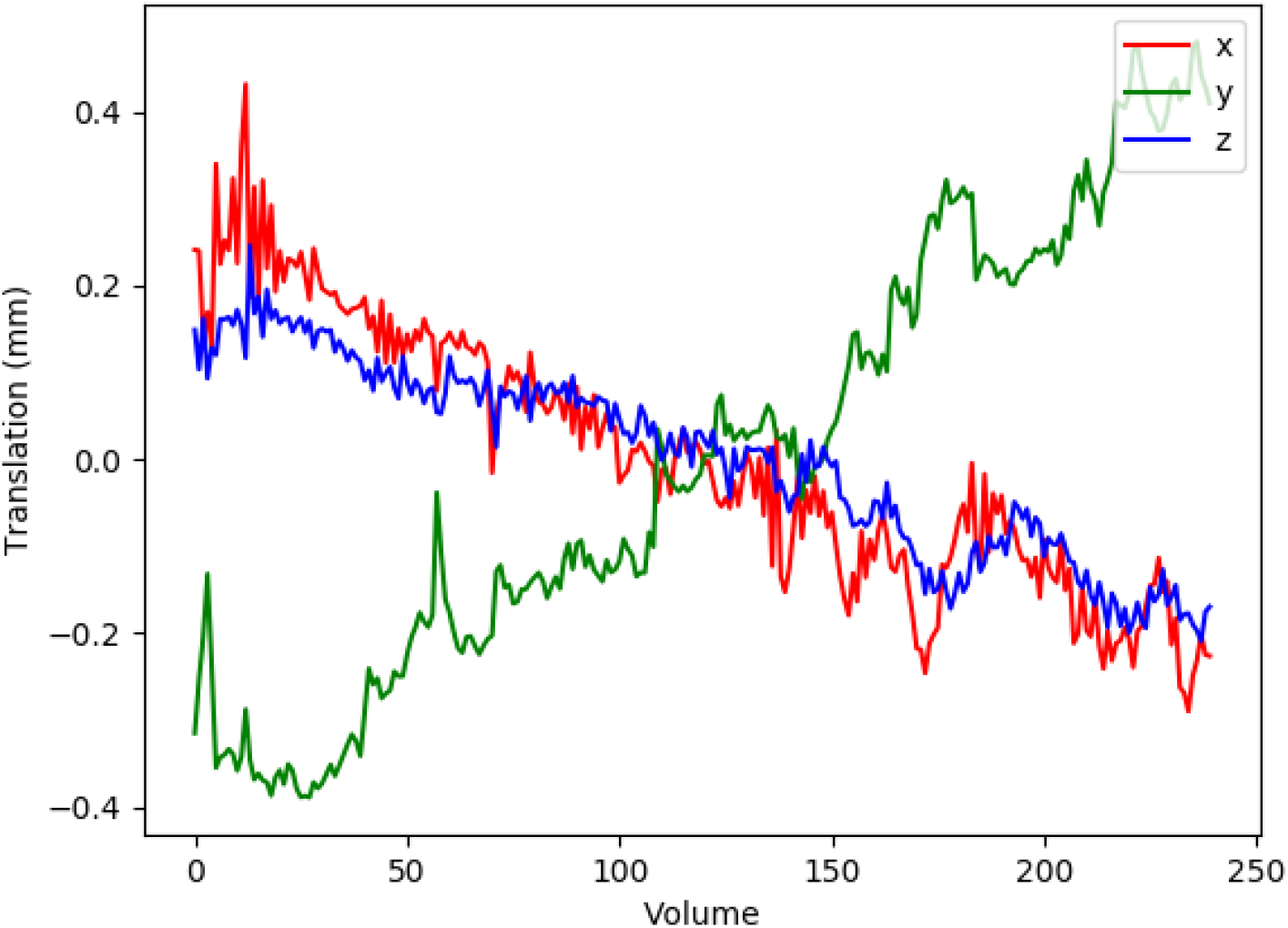
Motion Correction QA. Quality assurance figure from the `m2g-f` pipeline which plots the distance in the x, y, and z axes that each functional volume was moved during the motion correction using AFNI’s 3dvolreg.

